# A RESTful API to serve BAM file with OAuth2 compatible authorization

**DOI:** 10.1101/151787

**Authors:** Julien Delafontaine, Sylvain Pradervand

## Abstract

**Summary:** Bam-server is an open-source RESTful service to query slices of BAM files securely and manage their user accesses. A typical use case is the visualization of local read alignments in a web interface for variant calling diagnostic, without exposing sensitive data to unauthorized users through the network, and without moving the original - heavy - file. Bam-server follows the standard implementation of a protected resource server in the context of a typical token-based authorization protocol, supporting HMAC- and RSA-hashed signatures from an authorization server of choice.

**Availability:** The source code is available at <u>https://github.com/chuv-ssrc/bam-server-scala</u>, and a complete documentation can be found at <u>http://bam-server-scala.readthedocs.io/en/latest/</u>.

## 1 Introduction

The ongoing improvement in sequencing technologies generates larger and larger data footprints. Consequently, long-term data storage has to be managed to avoid unnecessary information duplication and to contain cost. Being able to access remotely large files is therefore of particular importance. In a typical whole genome sequencing application, sequencing reads are stored in files called BAM that contain the information about their alignment to a reference genome. BAM files are compressed, binary version of a standard text format called SAM (for Sequencing Alignment/Map) (Li *et al*. 2009). Accessing BAM files remains necessary even after genetic variants have been called, mainly because the visualization of local alignments gives a visual confirmation that compensates some weaknesses of variant callers.

In human genetics, BAM files are inherently sensitive and protecting their access is a fundamental requirement. While it is easy to serve open-access BAM files, controlling individual accesses of users to only their own data requires an additional layer, which can be tedious to write for IT specialists, and challenging for others. One of the most popular ways to handle data access authorizations securely is the OAuth2 scheme <u>(https://oauth.net/2/).</u> Thus, an OAuth2-compatible API has a real advantage when integrated with modern existing systems.

Bam-server provides such a secure authorization layer that is actually compatible with most token-based protocols, including OAuth2. Its REST API makes it usable as a web service; transferring only chunks of data through HTTP results in instant responses and thus a reactive user interface. With that in mind, it was also made to be directly usable in conjunction with IGV.js <u>(https://github.com/igvteam/igv.js),</u> a library embedding the popular Integrative Genomics Viewer (IGV) browser in Javascript in order to visualize read alignments directly in a web browser (Thorvaldsdóttir *et al*. 2013).

## 2 Method

One of the most popular authorization schemes is OAuth2, where the client application delegates the work of logging in the user to a remote authorization server. The client uses only the signed access “token” it gets in return from the authorization server to access a resource on behalf of that user. Bam-server represents the resource server in the authorization workflow (Fig. 1): having its own database of user permissions, it is able to verify signed tokens from registered applications to identify the requester and deny access when unauthorized. It leaves completely open the choice of both client application and authorization server. In particular, the authorization server should never know about BAM files, so all information concerning genetic samples can remain in the bam-server database.

**Fig. 1.**
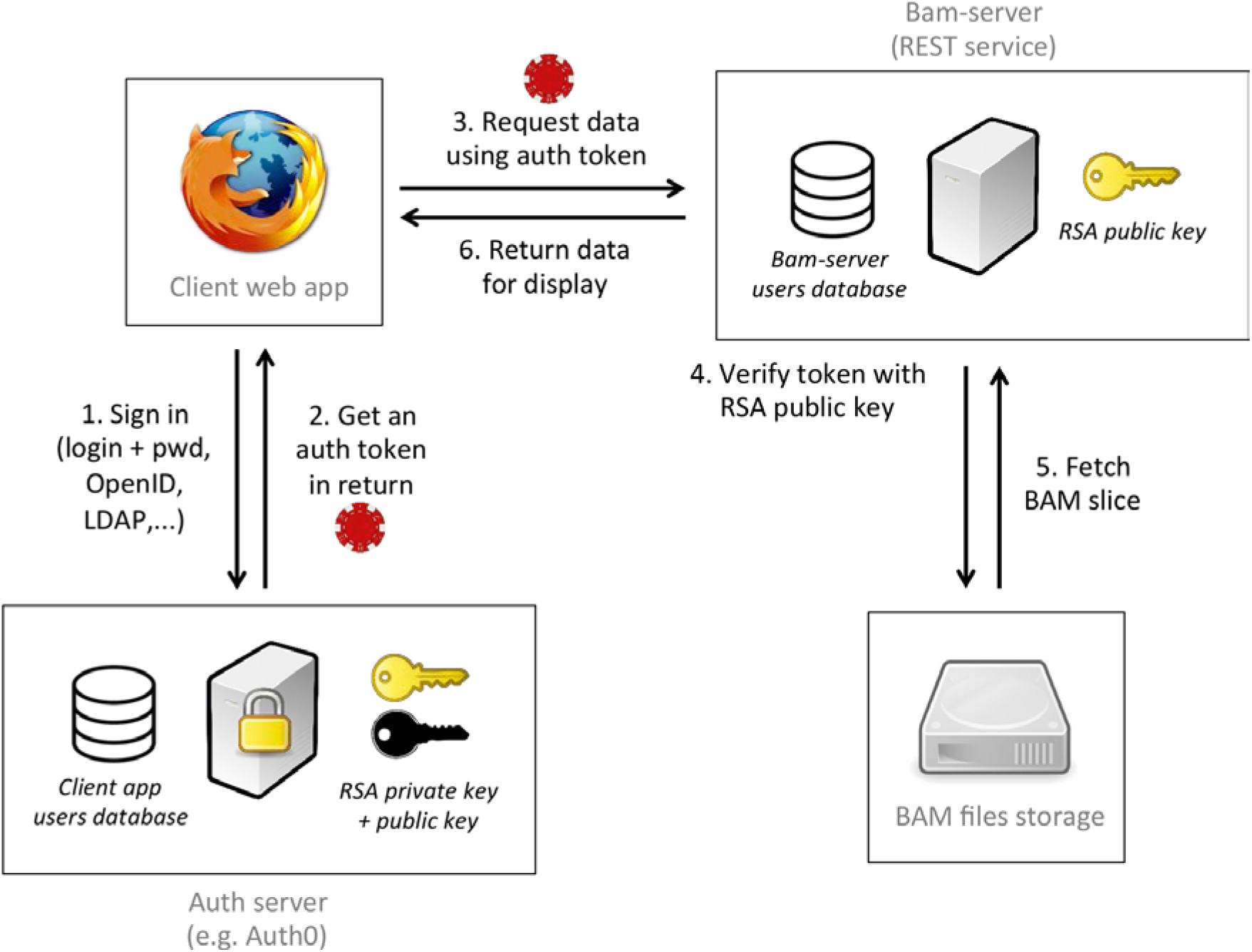
(1) The client app connects to the authorization (auth) server to request an auth token. (2) The auth server returns a signed JSON Web Token (JWT) that contains identifiers for the user and the client app. (3) The client requests a portion of a BAM file for a given sample from the bam-server using the REST API. (4) Bam-server verifies the token signature using a validation key, then checks the user and app identifiers against its own user database. (5) According to permissions, bam-server extracts the requested slice from the BAM file corresponding to the sample and (6) returns it to the client app.

## 3 Implementation

### 3.1. REST API

Bam-server provides a minimal RESTful API allowing to query portions of BAM files using three different strategies, depending on the use case: (1) extracting a range of bytes (based on BAM index, ex: IGV.js), (2) calling samtools (if available) to return a smaller BAM file for only one genomic region, or (3) using htsjdk (samtools Java API) to return reads in JSON format (Li *et al*. 2009). For instance,

~~~
GET http://bam-server.com/bam/json/sample1?region=chr1:222-333
~~~

will return a JSON array of all reads from “sample1” from genomic region “chr1:222-333” - only if the requester is allowed to access the corresponding BAM file.

Bam-server also provides a simple REST API for users and samples management.

### 3.2 Authorization

To verify permissions, all endpoints require to send with the HTTP request an Authorization header containing a signed JWT. Bam-server supports HMAC- and RSA-signed tokens. On each request, the signature is verified using either a shared secret key in the case of HMAC, or a public key in the case of RSA. Then the content is decoded and matched against the database that defines individual accesses.

### 3.3. Implementation example with IGV.js and Auth0

To illustrate how bam-server integrates with existing libraries and services, we created a demonstration client application using IGV.js for visualization and a dummy Auth0 account as authorization server. With these tools, it took only a few hours to build a good-looking page with a login screen and a reactive main window that permits to view read alignments in a web browser, and only for logged-in users that have been given access to the data.

The source code of the demo client is available in GitHub at https://github.com/chuv-ssrc/bam-server-client-demo

## 4 Discussion

Bam-server is a lightweight, easy-to-deploy service that assumes locally stored data and centralizes samples management and data access permissions under an administrator, which makes it best suited for small-scale deployment. It is tailored to allow research groups to quickly implement web clients that query parts of their data over a network without compromising the data. By returning either bytes, BAM or JSON, it adds flexibility on the client implementation. Another key advantage is its support for RSA signatures, so that it never has to store a secret.

Alternative solutions include NIH’s “bam-slicing” tool (<u>https://docs.gdc.cancer.gov/API/Users_Guide/BAM_Slicing/</u>), GA4GH Genomics API (<u>https://github.com/ga4gh/ga4gh-server</u>) or even a generic binary files server handling authorizations. They have different trade-offs in terms of scale, functionality, deployment effort and security. NIH’s “bam-slicing” is the closest software we could find in terms of functionality. Using a user-specific token valid for 30 days, in the form of an alphanumeric string hard-coded in a downloaded text file, it adds a security layer to its API to query slices of BAM files stored on NIH’s servers. We found that its source is not directly accessible or documented to use it apart from the Genomic Data Commons API interface.

